# Efficiency of genomic prediction of non-assessed single crosses

**DOI:** 10.1101/141440

**Authors:** José Marcelo Soriano Viana, Helcio Duarte Pereira, Gabriel Borges Mundim, Hans-Peter Piepho, Fabyano Fonseca e Silva

## Abstract

An important application of genomic selection in plant breeding is the prediction of untested single crosses (SCs). Most investigations on the prediction efficiency were based on tested SCs, using cross-validation. The main objective was to assess the prediction efficiency by correlating the predicted and true genotypic values of untested SCs (accuracy) and measuring the efficacy of identification of the best 300 untested SCs (coincidence), using simulated data. We assumed 10,000 SNPs, 400 QTLs, two groups of 70 selected DH lines, and 4,900 SCs. The heritabilities for the assessed SCs were 30, 60 and 100%. The scenarios included three sampling processes of DH lines, two sampling processes of SCs for testing, two SNP densities, DH lines from distinct and same populations, DH lines from populations with lower LD, two genetic models, three statistical models, and three statistical approaches. We derived a model for genomic prediction based on SNP average effects of substitution and dominance deviations. The prediction accuracy is not affected by the linkage phase. The prediction of untested SCs is very efficient. The accuracies and coincidences ranged from approximately 0.8 and 0.5, respectively, under low heritability, to 0.9 and 0.7, assuming high heritability. Additionally, we highlighted the relevance of the overall LD and evidenced that efficient prediction of untested SCs can be achieved for crops that show no heterotic pattern, for reduced training set size (10%), for SNP density of 1 cM, and for distinct sampling processes of DH lines, based on random choice of the SCs for testing.

## INTRODUCTION

Genomic selection is very commonly used in animal breeding programs, especially for dairy cattle (Van Eenennaam et al. 2014). The same cannot yet be said to the same degree concerning crop breeding. The main reasons for the effective application of genomic selection in livestock breeding are: it is efficient, that is, the process has high prediction accuracy, the cost of phenotyping (mainly progeny test) is higher than the cost of genotyping, and the process significantly shortens the selection cycle (Meuwissen et al. 2013). In spite of the many field- and simulation-based studies with genomic selection in plant breeding, in general the cost of phenotyping is often still much lower than the cost of genotyping, restricting its application in breeding programs. Jonas and de Koning (2013) consider that genomic selection has the potential to improve existing plant breeding schemes. However, based also on the high diversity and complexity of plant breeding methods, they stated that there are great obstacles to overcome.

An important application of genomic selection in plant breeding is the prediction of untested single crosses (genotypic value prediction) and testcrosses (general combining ability effect prediction) in hybrid breeding (Zhao et al. 2015). Genomic prediction of two- and three-way crosses has been investigated (Philipp et al. 2016). The prediction of untested single crosses was pioneered by Bernardo (1994), based on best linear unbiased prediction (BLUP). Many significant studies on prediction of untested single cross and testcross performance have been published in the last 23 years, focused on the assessment of the prediction accuracy. Most investigations were based on empirical data and estimated the prediction accuracy using a cross-validation procedure. Very few were based on simulated data (Li et al. 2017; Technow et al. 2012). With no exception, the inference was that prediction of untested single crosses and testcrosses can be an efficient, depending on heritability, training set size, and number of tested inbreds in hybrid combination (both, one, and none parents tested). Remarkably, this conclusion was drawn from studies differing in the type of molecular marker, density of markers, number of inbreds, level of relatedness, diversity, and linkage disequilibrium (LD) between inbreds, heterotic pattern, training set size, genetic model, and statistical approach (Zhao et al. 2015). Efficient prediction of barley two- and three-way crosses has been achieved when training and validation sets include the same class of hybrids (Philipp et al. 2016).

Most studies on genomic prediction of maize single cross performance published since 2011 have employed single nucleotide polymorphisms (SNP), with the number SNPs filtered ranging from 425 (Zhao et al. 2013a) to 39,627 (Technow et al. 2012). Based on the physical length of the maize genome (approximately 2,106 megabase pairs (Mb) according to Maize genetics and genomics database), the SNP density ranged from approximately 5 to 0.05 Mb, respectively. For grain yield, the relative prediction accuracies (computed as accuracy/root square of the heritability) in the two previously cited papers ranged from 0.27 to 0.62 and from 0.65 to 0.95, respectively. The number of inbreds in each heterotic group was highly variable too, ranging from six and nine (Bernardo 1994) to 75 and 75 (Technow et al. 2012). The relative accuracy observed by Bernardo (1994) ranged between 0.72 and 0.89. The number of testcrosses ranged between 255 (Windhausen et al. 2012) and 1,894 (Albrecht et al. 2014). The relative accuracies ranged from 0.46 to 0.52 and from 0.33 to 0.65, respectively. The level of relatedness ranged from non-related inbreds in each group (Technow et al. 2012) to a maximum average value of 0.58 (Bernardo 1995). The relative accuracy obtained by Bernardo (1995) ranged from 0.41 to 0.80. The common heterotic groups were Stiff Stalk and non-Stiff Stalk (Kadam et al. 1916) or Dent and Flint (Technow et al. 2014). The study of Bernardo (1996a) involved nine heterotic groups and the (statistically significant from zero) relative accuracies ranged from 0.43 to 0.88. No study provided clearly greater prediction accuracy of the additive-dominance model relative to the additive model. Finally, only with testcrosses the genomic BLUP (GBLUP) approach outperformed pedigree-based BLUP (Albrecht et al. 2014; Albrecht et al. 2011) concerning prediction accuracy.

Genomic prediction of single crosses has been made based on tested single crosses, using cross-validation. Thus, the estimated prediction accuracies are not for untested single crosses.
Consequently, none of the previous studies on efficiency of genomic prediction of single cross performance measured the efficacy of identification of the best untested single crosses. Our main objective was to assess the efficiency of prediction of untested single crosses by correlating the predicted and true genotypic values of untested single crosses (prediction accuracy) and measuring the efficacy of identification of the best 300 untested single crosses (coincidence index), using a large simulated data set. The secondary objectives were to highlight that the prediction accuracy depends primarily on the overall LD in the groups of selected doubled haploid (DH) lines, that the prediction efficiency when there is no heterotic pattern can be as high as the prediction efficiency when there are heterotic groups, and that the choice of single crosses for testing should be random, instead of selecting DH lines for a diallel, to maximize the prediction efficiency. Further, we derived a model for genomic prediction of untested single crosses based on the SNP average effects of substitution and dominance deviations.

## MATERIALS AND METHODS

### Theory

Generally, most papers on genomic selection presents only statistical aspects and the genetic models are deduced from gene to SNP effects. Importantly, when there is some quantitative genetics theory, the LD between QTLs and SNPs is usually completely ignored. The quantitative genetics theory developed in this paper provides a genetic model for genomic prediction of untested single crosses that accounts for the LD between QTLs and SNPs. The model developed offers the genetic background to the models fitted in important previously papers on prediction of untested single crosses and testcrosses (Massman et al. 2013; Technow et al. 2012; Albrecht et al. 2011).

#### LD in a group of selected DH or inbred lines

Consider a group of DH or inbred lines selected from a population or heterotic group. Assume also a QTL (alleles B/b) and a SNP (alleles C/c) where B and b are the alleles that increase and decrease the trait expression, respectively. Define the joint genotype probabilities as P(BBCC) = f_22_, P(BBcc) = f_20_, P(bbCC) = f_02_, and P(bbcc) = f_00_, where the subscript indicates the number of copies of the major allele (B and C). The measure of LD between the QTL and the SNP is Δ_bc_ = f_22_f_00_ – f_20_f_02_ (Kempthorne 1954) and the haplotype frequencies are P(BC) = f_22_ = p_b_p_c_ + Δ_bc_, P(Bc) = f_20_ = p_b_q_c_ – Δ_bc_, P(bC) = f_02_ = q_b_p_c_ – Δ_bc_, and P(bc) = f_00_ = q_b_q_c_ + Δ_bc_, where p is the frequency of the major allele (B or C) and q = 1 – p is the frequency of the minor allele (b or c). Notice that p_b_ = f_22_ + f_20_ and p_c_ = f_22_ + f_02_. It is important to highlight the fact that we are not assuming that the QTL and the SNP are linked and in LD in the population or heterotic group, because this is not a necessary condition for genomic prediction. But we are assuming that they are in LD in the group of DH or inbred lines. Furthermore, because of selection, genetic drift, and inbreeding (only for inbreds and linked QTLs and SNPs), the gene and genotypic frequencies and the LD values concerning the selected DH or inbred lines cannot be traced to the values in the population or heterotic group.

#### SNP genotypic values of DH or inbred lines

The average genotypic value for a group of selected DH or inbred lines is M_IL_ = m_b_ + (p_b_ – q_b_)a_b_, where m_b_ is the mean of the genotypic values of the homozygotes and a_b_ is the deviation between the genotypic value of the homozygote of higher expression and m_b_. Thus, the average SNP genotypic values for the DH or inbred lines CC and cc are

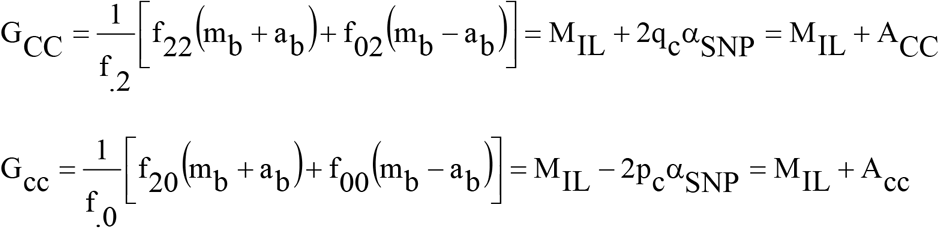

where 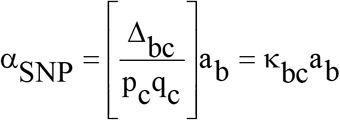 is the average effect of a SNP substitution in the group of DH or inbred lines and A is the SNP additive value for a DH or inbred line. Notice that E(A) = 0.

Assuming two QTLs (alleles B and b, and E and e) in LD with the SNP, the average effect of a SNP substitution in the selected DH or inbred lines is α_SNP_ = κ_bc_ a_b_ + κ_ce_ a_e_, where 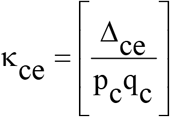. Thus, in general, the average effect of a SNP substitution (and the SNP additive value) is proportional to the LD measure and to the a deviation for each QTL that is in LD with the marker.

#### SNP genotypic values of single crosses

Aiming to maximize the heterosis, maize breeders commonly assess single crosses originating from selected DH or inbred lines from distinct heterotic groups. Consider n_1_ DH or inbred lines from a population or heterotic group and n_2_ DH or inbred lines from a distinct population or heterotic group. The average genotypic value for the single crosses derived by crossing the DH or inbred lines from group 1 with the DH or inbred lines from group 2 is

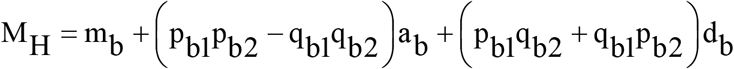

where d_b_ is the dominance deviation (the deviation between the genotypic value of the heterozygote and m_b_).

The average genotypic values for the single crosses derived from DH or inbred lines CC and cc of the group 1 are

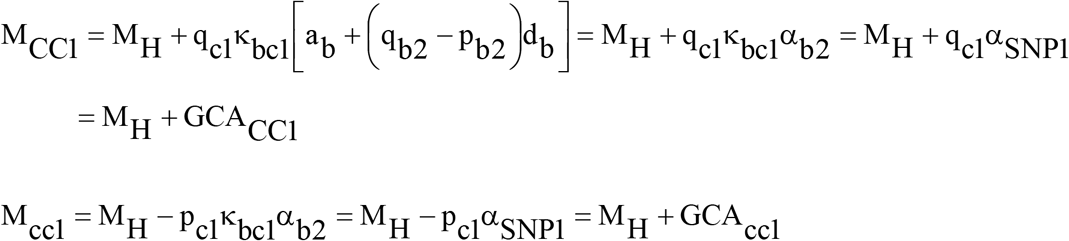

where α_b2_ is the average effect of allelic substitution in the population derived by random crosses between the DH or inbred lines of group 2, α_SNP1_ is the SNP effect of allelic substitution in the hybrid population relative to a SNP derived from group 1, and GCA stands for the general combining ability effect for a SNP locus. Notice that αSNP1 depends on the LD in group 1 (κ_bc1_ = Δ_bc1_/p_c1_q_c1_) and the average effect of allelic substitution in the population derived by random crosses between the DH or inbred lines of group 2. Further, E(GCA) = p_c1_GCA_CC1_ + q_c1_GCA_cc1_ = 0. Concerning the single crosses derived from DH or inbred lines CC and cc of the group 2 we have

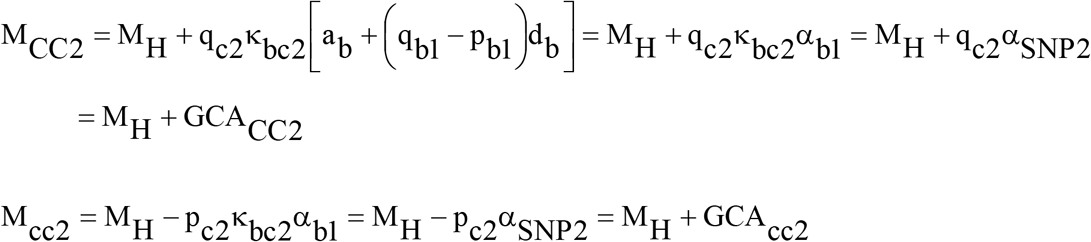

Notice that E(GCA) = 0 also. The average genotypic values for the single crosses concerning the SNP locus are

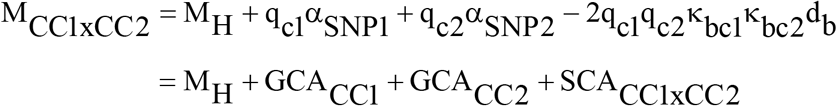

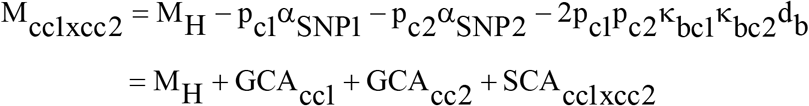

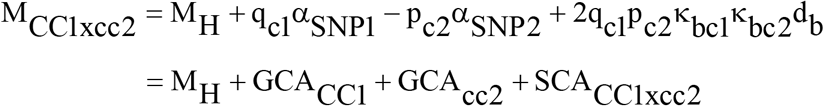

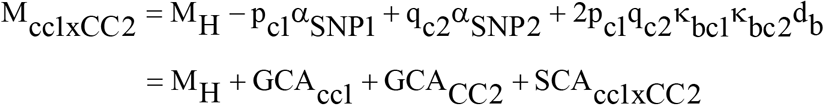

where κ_bc1_κ_bc2_d_b_ = d_SNP_ is the SNP dominance deviation in the hybrid population and SCA stands for the specific combining ability effect for a SNP locus. Notice that E(SCA) = p_c1_p_c2_SCA_CC1xCC2_ + p_c1_q_c2_SCA_CC1xcc2_ + q_c1_p_c2_SCA_cc1xCC2_ + q_c1_q_c2_SCA_cc1xcc2_ = 0 and, for each group, E(SCA|CC) = E(SCA|cc) = 0. That is, the expectation of the SNP SCA effects given a SNP genotype for the common DH or inbred line is also zero. Notice also that the four genotypic values depends on four unknown parameters (M_H_, α_SNP1_, α_SNP2_, and d_SNP_).

Assuming two QTLs (alleles B and b, and E and e) in LD with the SNP, the SNP dominance deviation is d_SNP_ = κ_bc1_κ_bc2_d_b_ + κ_ce1_κ_ce2_d_e_. Thus, generally, the SNP dominance deviation (and the SNP SCA effect) is proportional to the product of the LD values in both groups of DH or inbred lines and to the dominance deviation for each QTL that is in LD with the marker.

The previous model expressed as a function of the SNP GCA and SCA effects was proposed by Massman et al. (2013), but these authors assumed GCA_CC_ + GCA_cc_ = 0 (for each heterotic group and for each SNP) and SCA_CC1xCC2_ = SCA_cc1xcc2_ = −SCA_CC1xcc2_ = −SCA_cc1xCC2_. Technow et al. (2012) have used a standard extension from QTL to SNP, defining the single cross genotypic value for a SNP as a function of the SNP a and d deviations. That is, M = M_H_ + u_1_a_1_ + u_2_a_2_ + u_3_d, where u_1_ and u_2_ equal to 1/2 or −1/2 if the corresponding DH or inbred line is homozygous for distinct SNP alleles (CC or cc), and u3 equal to 0 if the single cross is homozygous or 1 if heterozygous.

#### SNP genotypic values of single crosses from DH or inbred lines derived from the same population or heterotic group

Well defined heterotic groups are known for maize, but not for special maize such as popcorn and sweet corn and for other crops such as wheat (Zhao et al. 2013b), rice (Xu et al. 2014), and barley (Philipp et al. 2016). Thus, for many breeders, it is interesting to know about the efficiency of genomic prediction of singles crosses when there are no heterotic groups. Assuming n DH or inbred lines derived from the same population or heterotic group, the average genotypic values for the single crosses concerning the SNP locus are

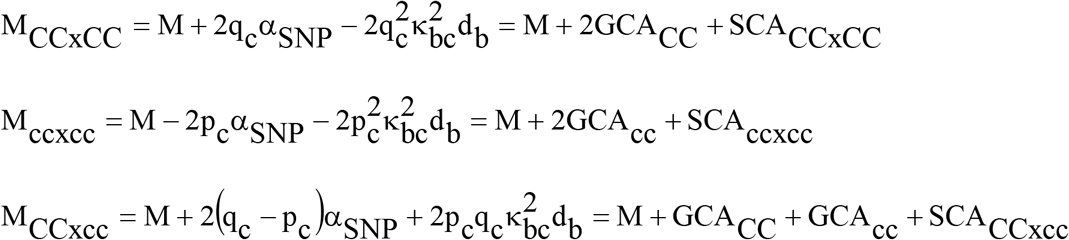

where M = m_b_ +(p_c_ – q_c_)a_b_ + 2p_c_q_c_d_b_ is the hybrid population mean, α_SNP_ = κ_bc_[a_b_ + (q_b_ – p_b_)d_b_] = κ_bc_α_b_ is the average effect of a SNP substitution in the hybrid population, and 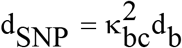 is the SNP dominance deviation. Notice that the SNP GCA effects are equal to half the SNP additive value for the single crosses (A), the SNP SCA effects are the SNP dominance deviations for the single crosses (D), and that the three genotypic values depends on three unknown parameters (M, α_SNP_, and d_SNP_). Notice also that E(GCA) = E(A) = E(SCA) = E(SCA|CC) = E(SCA|cc) = E(D) = 0.

#### Accuracy of single cross genomic prediction

Assuming a QTL and a SNP in LD in the two groups of DH or inbred lines, the predictor of the single cross QTL genotypic value is the single cross SNP genotypic value (because they are proportional). Thus, the covariance between the predictor and the genotypic value is

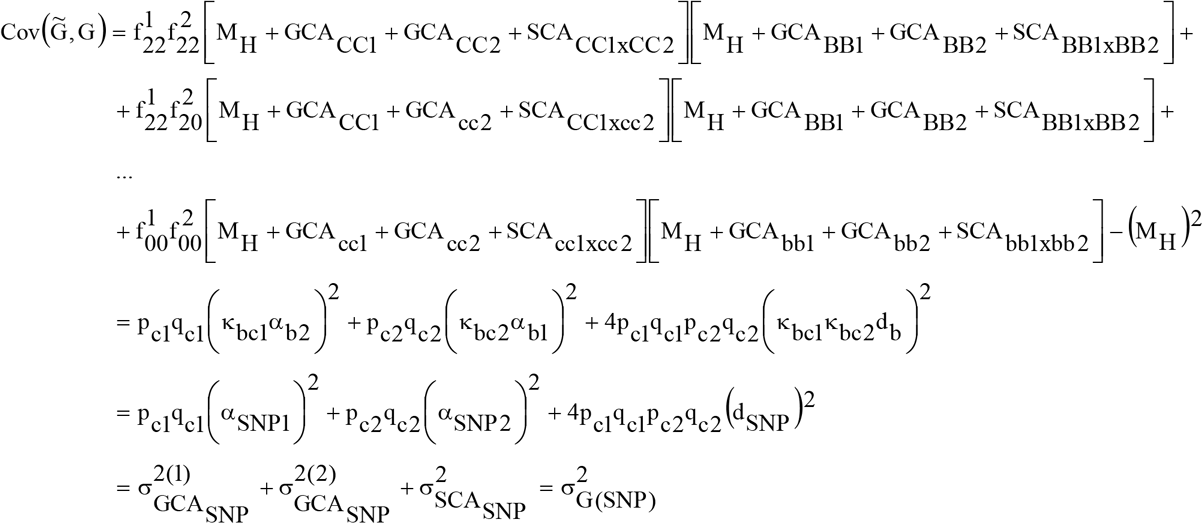

where the GCA and SCA effects for the QTL are GCA_BB1_ = q_b1_α_b2_, GCA_bb1_ = −p_b1_α_b2_, GCA_BB2_ = q_b2_α_b1_, GCA_bb2_ = −p_b2_α_b1_, SCA_BB1xBB2_ = −2q_b1_q_b2_d_b_, SCA_BB1xbb2_ = 2q_b1_p_b2_d_b_, SCA_bb1xBB2_ = 2p_b1_q_b2_d_b_, and SCA_bb1xbb2_ = −2p_b1_p_b2_d_b_, 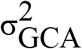 and 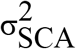 are the GCA and SCA variances for the SNP locus, and 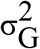 is the SNP genotypic variance. The GCA and SCA variances for the QTL are 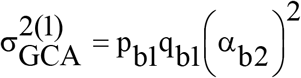, 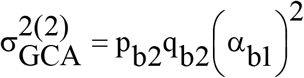, and 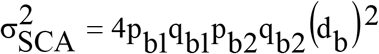. The QTL genotypic variance is 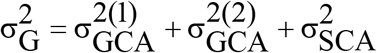 Thus, the single cross prediction accuracy is

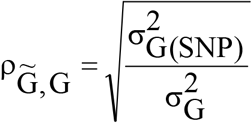

Assuming s SNPs,

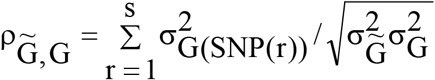

where 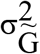 is the variance of the predicted single cross genotypic values and 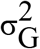 is the single cross genotypic variance. Further, 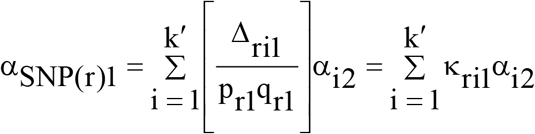, where k′ is the number of QTLs in LD with the SNP r) in group 1, and 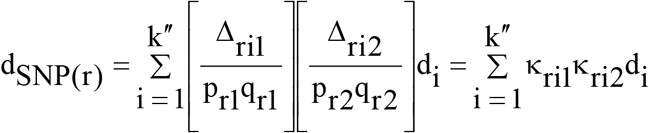 where k" is the number of QTLs in LD with the SNP r in both groups Notice that because the accuracy of genomic prediction of single crosses depends on the squares of the average effects of SNP substitution and the SNP dominance deviations, it is not affected by the linkage phase (coupling or repulsion), as it does not depend on linkage. But it depends on the magnitude of the LD in each group of DH or inbred lines.

Assuming single crosses derived from DH or inbred lines of a single population or heterotic group we have 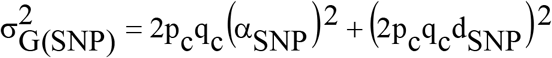 and 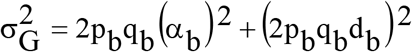. Therefore, the prediction accuracy of single crosses derived from DH or inbred lines from two distinct populations or heterotic groups differ from the prediction accuracy of single crosses resulting from DH or inbred lines obtained from each population or heterotic group.

### The statistical model for single cross genomic prediction

Assume n_1_ and n_2_ (several tens) DH or inbred lines from two populations or heterotic groups genotyped for s (thousands) SNPs and the experimental assessment of h (few hundred) single-crosses (h much lower than n_1_.n_2_) in e (several) environments (a combination of growing seasons, years, and locals). Defining y as the adjusted single cross phenotypic mean, the statistical model for prediction of the average effects of SNP substitution and the SNP dominance deviations is

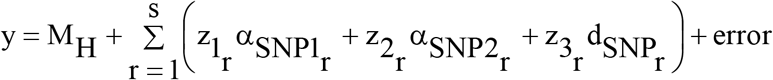

where z_1_r__ = q_r1_, z_2_r__ = q_r_2__, and z_3_r__ = −2q_r1_q_r2_ if the SNP genotypes for the two DH or inbred lines are CC (group 1) and CC (group 2), z_1_r__ = −p_r1_, z_2_r__ = −p_r2_, and z_3_r__ = −2p_r1_p_r2_ if the SNP genotypes for the DH or inbred lines are cc (group 1) and cc (group 2), z_1_r__ = q_r1_, z_2_r__ = −p_r2_, and z_3_r__ = 2q_r1_p_r2_ if the SNP genotypes for the DH or inbred lines are CC (group 1) and cc (group 2), and z_1_r__ = −p_r1_, z_2_r__ = q_r2_, and z_3_r__ = p_r1_q_r2_ if the SNP genotypes for the DH or inbred lines are cc (group 1) and CC (group 2).

Regarding the single crosses obtained from DH or inbred lines of the same population or heterotic group we have

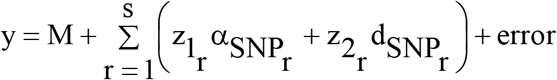

where z_1_r__ = 2q_r_ and 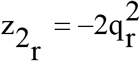 if the SNP genotypes for the two crossed DH or inbred lines are CC and CC, z_1_r__ = −2p_r_ and 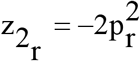 if the SNP genotypes for the two DH or inbred lines are cc and cc, and z_1_r__ = 2(q_r_ – p_r_) and z_2_r__ = 2p_r_q_r_ if the SNP genotypes for the DH or inbred lines are CC and cc.

The statistical problem of genomic prediction when there are a very large number of molecular markers and relatively few observations have been addressed thorough several regularized whole-genome regression and prediction methods (Daetwyler et al. 2013; de Los Campos et al. 2013). Based on one of these approaches, the SNP average effects of substitution and SNP dominance deviations are predicted and used to provide genomic prediction of non-assessed single crosses. The predicted genotypic value for a non-assessed single cross of DH or inbred lines from two groups is

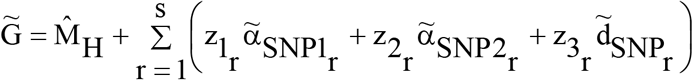

For a non-assessed single cross of DH or inbred lines from the same group, the predicted genotypic value is

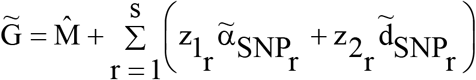

### Simulation

The SNP and QTL genotypic data for DH lines, the QTL genotypic data of single crosses, and the phenotypic data for DH lines and single crosses were simulated using the software *REALbreeding.* The program has been developed by the first author using the software *REALbasic 2009* (Viana et al. 2017a; Viana et al. 2017b; Viana et al. 2016; Azevedo et al. 2015; Viana et al. 2013). Based on our input, the software distributed 10,000 SNPs and 400 QTLs in ten chromosomes (1,000 SNPs and 40 QTLs by chromosome). The average SNP density was 0.1 cM. The QTLs were distributed in the regions covered by the SNPs (approximately 100 cM/chromosome). Initially, *REALbreeding* sampled 700 DH lines from two non-inbred populations (heterotic groups) in LD (350 from each population). The populations were composites of two populations in linkage equilibrium. In a composite, there is LD only for linked SNPs and QTLs (Viana et al. 2016). The number of DH lines from each S_0_ plant was one (scenario 1) or ranged from 1 to 5 (scenario 2). We also sampled 350 DH lines from each population after three generations of selfing (using a single seed descent process). The number of DH lines from each S_3_ plant ranged from 1 to 5 (scenario 3). For each scenario, the software then crossed 70 selected DH lines from each population, using a diallel design. The heritability for the DH lines was 30%.

The genotypic values of the DH lines and of the single crosses were generated assuming a single set of 400 QTLs and two degrees of dominance. To simulate grain yield and expansion volume, a measure of popcorn quality, we defined positive dominance (0 < (d/a)_i_ ≤ 1.2, i = 1, &, 400) and bidirectional dominance (−1.2 ≤ (d/a)_i_ ≤ 1.2), respectively, where d/a is the degree of dominance. To compute the genotypic values, *REALbreeding* used our input relative to the maximum and minimum genotypic values for homozygotes. For grain yield and expansion volume, we defined 140 and 30 g/plant and 55 and 15 mL/g, respectively. The phenotypic values were obtained from the sum of the population mean, genotypic value, and experimental error. The error variance was computed from the broad sense heritability. To avoid outliers, we defined the maximum and minimum phenotypic values as 160 and 10 g/plant and 65 and 5 mL/g.

The heritabilities for the assessed single crosses were 30, 60, and 100%. Thus, the genotypic value prediction accuracies of the assessed single crosses were 0.55, 0.77, and 1.00, respectively. For each scenario were processed 50 resamplings of 30 and 10% of the single crosses (1,470 and 490 assessed single crosses). That is, we predicted 70 and 90% of the single crosses (3,430 and 4,410 non-assessed single crosses). Additionally, to assess the relevance of the number of DH lines sampled, we fixed the number of DH lines to achieve the same number of assessed single crosses, using a diallel. That is, we sampled 50 times 38 and 22 DH lines in each group for a diallel (scenario 4), generating 1,444 and 484 single crosses for assessment, respectively. We denote these processes as sampling of single crosses (scenarios 1 to 3) and sampling of DH lines (scenario 4). Other additional scenarios were: genomic prediction of single crosses from selected DH lines from same heterotic group (interestingly for wheat, rice, and barley breeders, for example) (scenario 5) and from selected DH lines from populations with lower LD (scenario 6), to emphasize that the prediction accuracy depends on the LD in the groups of DH or inbred lines. A last scenario (seventh) was genomic prediction of single crosses under an average density of one SNP each cM. This lower density was obtained by random sampling of 100 SNPs per chromosome using a *REALbreeding* tool (*sampler*). To investigate the single cross prediction efficiency based on our model and on the models proposed by Massman et al. (2013) and Technow et al. (2012), we used another *REALbreeding* tool (*Incidence matrix*) to generate the incidence matrices for the three models and for the two DH lines sampling processes. To assess the relevance of the SCA effects prediction on genomic prediction of single cross performance, we also fitted the additive model (including only the GCA effects). For comparison purpose, we also processed single cross prediction based on GBLUP (with the observed additive and dominance relationship matrices) and pedigree-based BLUP (with the expected additive and dominance relationship matrices).

### Statistical analysis

The methods used for prediction were ridge regression BLUP (RR-BLUP), GBLUP and BLUP. For the analyses we used the *rrBLUP* package (Endelman 2011). The accuracies of single cross genotypic value prediction were obtained by the correlation between the true values of the non-assessed single crosses computed by *REALbreeding* and the values predicted by RR-BLUP, GBLUP, and BLUP. We also computed the efficiency of identification of the 300 non-assessed single crosses of higher genotypic value (coincidence index). The coincidence index was computed from the 300 higher predicted untested single crosses as the number of predicted untested single crosses among the 300 untested single crosses of greater true genotypic value/300. For each DH lines derivation process and heritability, the parametric average coincidence index was computed from the average phenotypic values of the 4,900 single crosses as the number of single crosses among the 300 single crosses of greater true genotypic value/300. Regarding grain yield, for heritability of 30% the coincidence index was 0.2533, 0.2833, and 0.2433 assuming one DH line per S_0_ plant, one to five DH lines per S_0_ plant, and one to five DH lines per S_3_ plant, respectively. The corresponding values for heritability of 60% were, respectively, 0.4800, 0.4900, and 0.4567. Concerning expansion volume, the corresponding values for heritabilities of 30 and 60% were, respectively, 0.2600, 0.2833, and 0.2700, and 0.4733, 0.5100, and 0.4533. The assumed average parametric coefficient index was 0.26 and 0.48 for heritabilities of 30 and 60%, respectively, for both traits. For the population structure analysis we employed *Structure* (Falush et al. 2003) and fitted the no admixture model with independent allelic frequencies. The number of SNPs, sample size, burn-in period, and number of MCMC (Markov chain Monte Carlo) replications were 1,000 (sampled at random), 140 (70 DH lines from each population), 10,000, and 40,000, respectively. The number of populations assumed (K) ranged from 1 to 4, and the most probable *K* value was determined based on the inferred plateau method (Viana et al. 2013). The LD analyses were performed with *Haploview* (Barrett et al. 2005).

### Data availability

*REALbreeding* is available upon request. The data set is available at https://doi.org/10.6084/m9.figshare.5035130.v3. Data citation: Viana, José Marcelo Soriano; Pereira, Helcio Duarte; Mundim, Gabriel Borges; Piepho, Hans-Peter; Fonseca e Silva, Fabyano (2017): Efficiency of genomic prediction of non-assessed single crosses. figshare. https://doi.org/10.6084/m9.figshare.5035130.v3

## RESULTS

The parametric mean and genotypic variance in the populations 1 and 2 were 108.5 and 87.3 (g/plant) and 4.7680 and 6.2580 (g/plant)^2^, respectivelly. The DH lines derivation processes (one and one to five per S_0_ plant and one to five per S_3_ plant) provided, for each population, selected DH lines with similar mean (approximately 97 and 76 g/plant for populations 1 and 2), inbreeding depression (approximately −10 and −13% for populations 1 and 2), and genotypic variance (approximately 6 and 7 (g/plant)^2^ for populations 1 and 2) and groups of single crosses also similar for mean (approximately 103 g/plant), heterosis (approximately 19%), and genotypic variance (approximately 4 (g/plant)^2^). Because we derived one to few DH lines from unrelated S_0_ and S_3_ plants, the average level of relatedness between the selected DH lines was very low (zero and zero, 0.0041 and 0.0041, and 0.0054 and 0.0074 assuming one DH line per S0, one to five DH lines per S_0_, and one to five DH lines per S_3_, for populations 1 and 2, respectively). Concerning SNP data, the frequency distribution of the minor allele frequency (MAF) and the absolute value of the difference between a SNP allele frequency were also similar for both groups of selected DH lines, regardless of the DH line derivation process (Figure 1a, b, c). The average MAF was 0.33, regardless of the population and DH line derivation process. However, the evidence obtained by the population structure analysis was that the DH lines belong to two distinct subpopulations (suggested *K* equal to 2.4 by the inferred plateau method). The percentages of non-polymorphic SNPs were very low (0.1 to 0.4%). No differences between allelic frequencies were observed for only 1.7 to 2.1% of the SNPs. For approximately 70% of the SNPs, the absolute difference between allelic frequencies ranged from 0.1 to 0.6. Regarding LD, for the groups of selected DH lines the evidence based on the analysis of chromosome 1 (no difference between chromosomes is expected) is that LD extents for up to 35 cM, regardless of the DH lines derivation process (Figure 1c, d). Ignoring the non-significant LD values (LOD score lower than 3), for 17 to 20% of the SNP pairs the r^2^ values ranged from 0.2 to 0.5 (average of 0.16, regardless of the DH lines group and derivation process).

**Figure 1.**
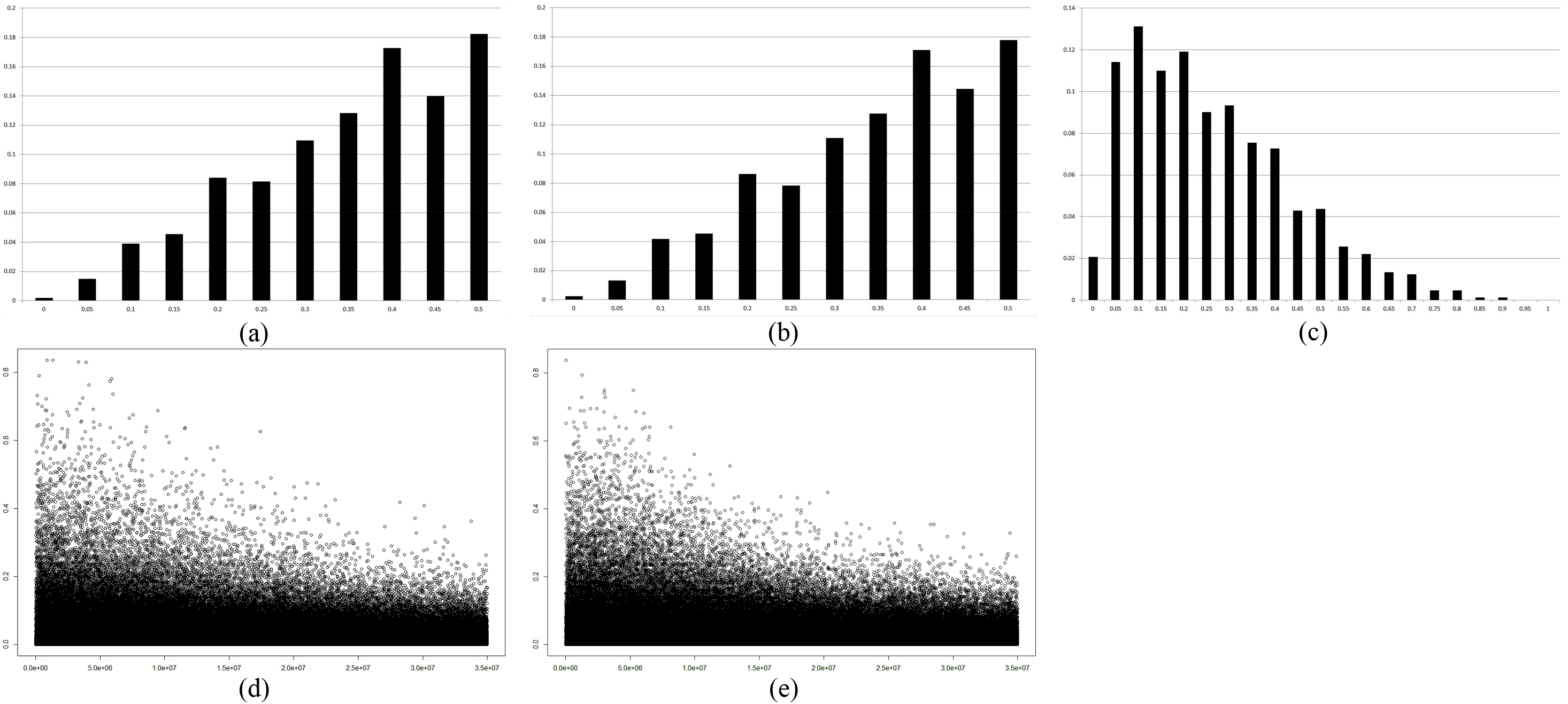
Frequency distribution of the MAF in the groups of selected DH lines (a and b) and the absolute value of the difference between a SNP allele frequency (c), and LD (r^2^) in relation to distance (cM) in the two groups of selected DH lines (d and e), regarding SNPs in chromosome 1 separated by zero to 35 cM, assuming one DH line per S_0_ plant.

Assuming our model, average SNP density of 0.1 cM, training set size of 30%, positive dominance (grain yield), additive-dominance model, and sampling of single crosses, the prediction accuracies of the non-assessed single crosses were greater than the accuracies of the assessed single crosses for low (up to 46% higher) and intermediate (up to 16% higher) heritabilities (Table 1; Figure 2a). As the prediction accuracy of assessed single crosses approaches 1.0, the accuracy of the non-assessed single crosses approaches approximately 0.9 (up to 11% lower). Sampling one to five DH lines per S3 plant was only slightly superior to the other DH lines derivation processes, regardless of the prediction accuracy of the assessed single crosses (up to 5% higher). Fitting the additive model provided essentially the same prediction accuracies since the maximum decrease was approximately 1%. No significant differences between the prediction accuracies of non-assessed single crosses were also observed assuming bidirectional dominance (expansion volume).

**Figure 2.**
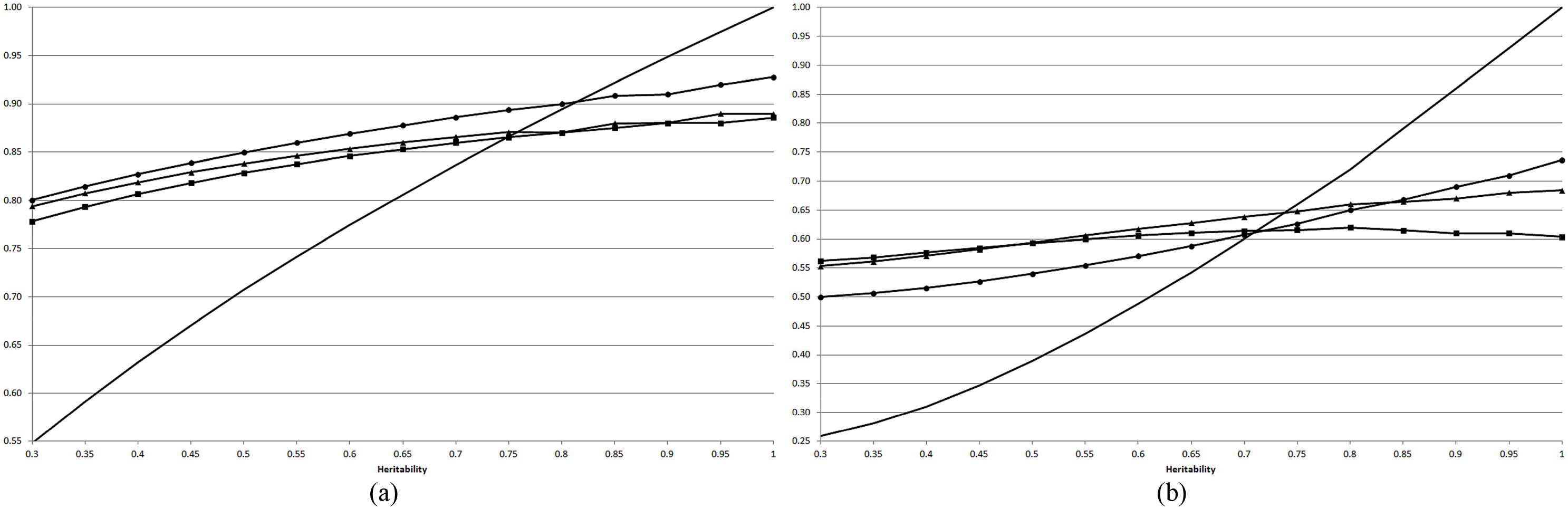
Predicted accuracies (a) and coincidence indexes (b) for untested single crosses (square: 1/S_0_; triangle: 1–5/S_0_; circle: 1–5/S_3_), and parametric accuracies and coincidence indexes for tested single crosses (continuous line), assuming our model, average SNP density of 0.1 cM, training set size of 30%, positive dominance (grain yield), additive-dominance model, and sampling of single crosses.

**Table 1.** Average prediction accuracies of non-assessed single crosses and its standard deviation, assuming single crosses from selected DH lines, 30 and 10% of assessed single crosses, two traits (grain yield - GY, g/plant, and expansion volume - EV, mL/g), two sampling processes of single crosses, four statistical models, three DH lines sampling processes, two genetic models, and three accuracies of assessed single crosses

| Trait | Samp. proc. | Statistical model | DH lines | Gen. mod. | Accuracy of assessed single crosses |  |  |
| --- | --- | --- | --- | --- | --- | --- | --- |
|  |  |  |  |  | 0.55 | 0.77 | 1.00 |
| GY | SCs | Viana et al. | 1/S<sub>0</sub> | AD | 0.7790 ± 0.0124 | 0.8447 ± 0.0066 | 0.8859 ± 0.0018 |
|  |  |  |  | A | 0.7688 ± 0.0132 | 0.8380 ± 0.0067 | 0.8821 ± 0.0019 |
|  |  |  | 1-5/S<sub>0</sub> | AD | 0.7947 ± 0.0125 | 0.8525 ± 0.0072 | 0.8896 ± 0.0025 |
|  |  |  |  | A | 0.7895 ± 0.0126 | 0.8465 ± 0.0077 | 0.8858 ± 0.0027 |
|  |  |  | 1-5/S<sub>3</sub> | AD | 0.8010 ± 0.0145 | 0.8678 ± 0.0054 | 0.9276 ± 0.0025 |
|  |  |  |  | A | 0.7954 ± 0.0145 | 0.8627 ± 0.0056 | 0.9238 ± 0.0026 |
|  |  |  | 1-5/S<sub>3</sub> | AD<sup>a</sup> | 0.7718 ± 0.0161 | 0.8371 ± 0.0079 | 0.8888 ± 0.0043 |
|  |  |  | 1-5/S<sub>3</sub> | AD<sup>b</sup> | 0.6836 ± 0.0277 | 0.7885 ± 0.0139 | 0.8817 ± 0.0049 |
|  |  |  | 1/S<sub>0</sub> | AD<sup>c</sup> | 0.8293 ± 0.0131 | 0.8944 ± 0.0049 | 0.9479 ± 0.0017 |
|  |  |  | 1-5/S<sub>3</sub> | AD<sup>d</sup> | 0.8267 ± 0.0082 | 0.8928 ± 0.0043 | 0.9083 ± 0.0023 |
|  |  | Massman et. al.<sup>e</sup> | 1/S<sub>0</sub> | AD | 0.7874 ± 0.0118 | 0.8519 ± 0.0053 | 0.8924 ± 0.0026 |
|  |  |  | 1-5/S<sub>0</sub> | AD | 0.7982 ± 0.0140 | 0.8622 ± 0.0055 | 0.8973 ± 0.0025 |
|  |  |  | 1-5/S<sub>3</sub> | AD | 0.8074 ± 0.0112 | 0.8753 ± 0.0056 | 0.9314 ± 0.0026 |
|  |  | GBLUP | 1/S<sub>0</sub> | AD | 0.7841 ± 0.0122 | 0.8477 ± 0.0064 | 0.8906 ± 0.0019 |
|  |  |  | 1-5/S<sub>0</sub> | AD | 0.7973 ± 0.0124 | 0.8574 ± 0.0070 | 0.8978 ± 0.0019 |
|  |  |  | 1-5/S<sub>3</sub> | AD | 0.7911 ± 0.0146 | 0.8639 ± 0.0056 | 0.9319 ± 0.0023 |
|  |  | BLUP | 1/S<sub>0</sub> | AD | 0.7855 ± 0.0129 | 0.8541 ± 0.0059 | 0.8899 ± 0.0019 |
|  |  |  | 1-5/S<sub>0</sub> | AD | 0.7803 ± 0.0143 | 0.8435 ± 0.0074 | 0.8830 ± 0.0024 |
|  |  |  | 1-5/S<sub>3</sub> | AD | 0.7227 ± 0.0203 | 0.7915 ± 0.0077 | 0.8373 ± 0.0048 |
|  | DHs | Viana et al. | 1/S<sub>0</sub> | AD | 0.5012 ± 0.0416 | 0.5117 ± 0.0467 | 0.5343 ± 0.0467 |
|  |  |  | 1-5/S<sub>0</sub> | AD | 0.4827 ± 0.0423 | 0.5000 ± 0.0420 | 0.5036 ± 0.0465 |
|  |  |  | 1-5/S<sub>3</sub> | AD | 0.5799 ± 0.0437 | 0.6106 ± 0.0413 | 0.6357 ± 0.0429 |
| EV | SCs | Viana et al. | 1/S<sub>0</sub> | AD | 0.7779 ± 0.0157 | 0.8458 ± 0.0069 | 0.8820 ± 0.0024 |
|  |  |  | 1-5/S<sub>0</sub> | AD | 0.8019 ± 0.0155 | 0.8656 ± 0.0050 | 0.9055 ± 0.0020 |
|  |  |  | 1-5/S<sub>3</sub> | AD | 0.7589 ± 0.0143 | 0.8424 ± 0.0058 | 0.9165 ± 0.0027 |

<sup>a</sup>density of 1 cM; <sup>b</sup>training set of 490 single crosses (10%); <sup>c</sup>after 10 generations of random crosses; <sup>d</sup>single crosses from DH lines of the same population; <sup>e</sup>and Technow et al..

The differences compared to positive dominance ranged from approximately −5 to 2%. However, a striking difference was observed between the sampling processes of single crosses for testing. Random sampling of single crosses provided higher prediction accuracies of non-assessed single crosses, compared to sampling DH lines for a diallel. The increases in the accuracies by sampling single crosses ranged from approximately 38 to 77%, proportional to the heritability. Decreasing the average SNP density to 1 cM led to a slight decrease in the prediction accuracy of non-assessed single crosses of approximately −4%). Decreasing the training set size to 10% decreased the prediction accuracy of non-assessed single crosses in approximately −5 to −15%, inversely proportional to the heritability. To establish that the prediction accuracy of non-assessed single crosses depends on the level of (overall) LD in the groups of selected DH or inbred lines, we derived DH lines from the same base populations after 10 generations of random crosses (to decrease the LD). The accuracies were also high, ranging from 0.83 to 0.95, proportional to the heritability. The prediction accuracies of non-assessed single crosses from DH lines of the same population were equivalent to the accuracies for single crosses derived from DH lines belonging to distinct heterotic groups, ranging from 0.83 to 0.91, also proportional to the heritability. Comparing our statistical model with the models proposed by Massman et al. (2013) and Technow et al. (2012), we observed no differences for the prediction accuracies of non-assessed single crosses (maximum difference of 1%). Interestingly, the Massman et al. (2013) and Technow et al. (2012) models provide identical accuracies. Finally, no significant differences between the prediction accuracies for RR-BLUP, GBLUP, and BLUP occurred (maximum of 2%), excepting for one to five DH lines per S_3_ plant, where BLUP was 9 to 10% inferior, regardless of the heritability.

Concerning the coincidence index, in general the inferences are the same established from the prediction accuracy analysis (Table 2; Figure 2b). There were no differences between the coincidence indexes regarding our model and the models proposed by Massman et al. (2013) and Technow et al. (2012) (maximum difference of 3%), and between the RR-BLUP, GBLUP, and BLUP approaches, except for one to five DH lines per S_3_ plant, where BLUP was −19 to −27% inferior, proportional to the heritability. The coincidence indexes were also high for single crosses derived from selected DH lines obtained from the base populations with lower LD (ranging from 0.55 to 0.76, proportional to the heritability) and from selected DH lines of the same population (ranging from 0.61 to 0.76, also proportional to the heritability). Sampling single crosses for assessment also provided higher coincidence index compared to sampling DH lines for a diallel (39 to 98% higher, proportional to the heritability). Decreasing the SNP density and the training set size decreased the coincidence index from 5 to 10% (proportional to the heritability) and from 17 to 26% (inversely proportional to the heritability), respectively. The maximum difference in the coincidence index by fitting the additive-dominant and the additive models was −3%. Only for one DH line per S_0_ plant the coincidence indexes assuming bidirectional dominance were slightly greater than the values assuming positive dominance (9 to 14% greater). This sampling process of DH lines provided the higher values of coincidence index, compared to the other sampling processes (7 to 26% higher, inversely proportional to the heritability). Finally, the coincidence index of the non-assessed single crosses are greater than the parametric values for all assessed single crosses assuming low (up to 117% higher) and intermediate (up to 39% higher) heritabilities (Table 1). However, as the parametric coincidence of assessed single crosses approaches 1.0, the coincidence values of the non-assessed single crosses approach approximately 0.60 to 0.74 (up to 26 to 40% lower), depending on the DH line sampling process.

**Table 2.** Average coincidence of the best 300 predicted single crosses and its standard deviation, assuming single crosses from selected DH lines, 30 and 10% of assessed single crosses, two traits (grain yield - GY, g/plant, and expansion volume - EV, mL/g), two sampling processes of single crosses, four statistical models, three DH lines sampling processes, two genetic models, and three parametric coincidence of assessed single crosses

| Trait | Samp. proc. | Statistical model | DH lines | Gen. mod. | Coincidence of assessed single crosses |  |  |
| --- | --- | --- | --- | --- | --- | --- | --- |
|  |  |  |  |  | 0.26 | 0.48 | 1.00 |
| GY | SCs | Viana et al. | 1/S<sub>0</sub> | AD | 0.4523 ± 0.0334 | 0.5525 ± 0.0190 | 0.6037 ± 0.0170 |
|  |  |  |  | A | 0.4396 ± 0.0346 | 0.5449 ± 0.0176 | 0.5976 ± 0.0172 |
|  |  |  | 1-5/S<sub>0</sub> | AD | 0.5686 ± 0.0273 | 0.6369 ± 0.0221 | 0.6842 ± 0.0140 |
|  |  |  |  | A | 0.5640 ± 0.0283 | 0.6299 ± 0.0221 | 0.6816 ± 0.0152 |
|  |  |  | 1-5/S<sub>3</sub> | AD | 0.5129 ± 0.0235 | 0.6044 ± 0.0200 | 0.7363 ± 0.0183 |
|  |  |  |  | A | 0.5063 ± 0.0225 | 0.5993 ± 0.0193 | 0.7305 ± 0.0190 |
|  |  |  | 1-5/S<sub>3</sub> | AD<sup>a</sup> | 0.4881 ± 0.0278 | 0.5691 ± 0.0229 | 0.6620 ± 0.0215 |
|  |  |  | 1-5/S<sub>3</sub> | AD<sup>b</sup> | 0.3805 ± 0.0511 | 0.4797 ± 0.0354 | 0.6087 ± 0.0233 |
|  |  |  | 1/S<sub>0</sub> | AD<sup>c</sup> | 0.5528 ± 0.0298 | 0.6489 ± 0.0203 | 0.7571 ± 0.0162 |
|  |  |  | 1-5/S<sub>3</sub> | AD<sup>d</sup> | 0.6116 ± 0.0214 | 0.7156 ± 0.0150 | 0.7581 ± 0.0166 |
|  |  | Massman et. al.<sup>e</sup> | 1/S<sub>0</sub> | AD | 0.4670 ± 0.0346 | 0.5663 ± 0.0174 | 0.6157 ± 0.0157 |
|  |  |  | 1-5/S<sub>0</sub> | AD | 0.5651 ± 0.0310 | 0.6431 ± 0.0164 | 0.6955 ± 0.0144 |
|  |  |  | 1-5/S<sub>3</sub> | AD | 0.5279 ± 0.0291 | 0.6139 ± 0.0204 | 0.7423 ± 0.0172 |
|  |  | GBLUP | 1/S<sub>0</sub> | AD | 0.4622 ± 0.0308 | 0.5660 ± 0.0190 | 0.6092 ± 0.0163 |
|  |  |  | 1-5/S<sub>0</sub> | AD | 0.5650 ± 0.0280 | 0.6384 ± 0.0204 | 0.6849 ± 0.0137 |
|  |  |  | 1-5/S<sub>3</sub> | AD | 0.5010 ± 0.0245 | 0.5937 ± 0.0216 | 0.7294 ± 0.0168 |
|  |  | BLUP | 1/S<sub>0</sub> | AD | 0.4641 ± 0.0331 | 0.5709 ± 0.0176 | 0.6081 ± 0.0127 |
|  |  |  | 1-5/S<sub>0</sub> | AD | 0.5531 ± 0.0323 | 0.6272 ± 0.0194 | 0.6699 ± 0.0130 |
|  |  |  | 1-5/S<sub>3</sub> | AD | 0.4172 ± 0.0258 | 0.4731 ± 0.0211 | 0.5377 ± 0.0196 |
|  | DHs | Viana et al. | 1/S<sub>0</sub> | AD | 0.2753 ± 0.0374 | 0.3056 ± 0.0445 | 0.3169 ± 0.0401 |
|  |  |  | 1-5/S<sub>0</sub> | AD | 0.3268 ± 0.0642 | 0.3400 ± 0.0691 | 0.3461 ± 0.0728 |
|  |  |  | 1-5/S<sub>3</sub> | AD | 0.3699 ± 0.0583 | 0.3931 ± 0.0579 | 0.4300 ± 0.0633 |
| EV | SCs | Viana et al. | 1/S<sub>0</sub> | AD | 0.5156 ± 0.0331 | 0.6081 ± 0.0159 | 0.6599 ± 0.0146 |
|  |  |  | 1-5/S<sub>0</sub> | AD | 0.5506 ± 0.0285 | 0.6337 ± 0.0203 | 0.6944 ± 0.0141 |
|  |  |  | 1-5/S<sub>3</sub> | AD | 0.4746 ± 0.0294 | 0.5843 ± 0.0174 | 0.7141 ± 0.0171 |

<sup>a</sup>density of 1 cM; <sup>b</sup>training set of 490 single crosses (10%); <sup>c</sup>after 10 generations of random crosses; <sup>d</sup>single crosses from DH lines of the same population; <sup>e</sup>and Technow et al..

## DISCUSSION

Twenty-three years ago, Bernardo (1994) first suggested to use BLUP for predicting untested maize single cross performance. Based on the prediction accuracies obtained by Bernardo (1994, 1995, 1996a, 1996b, 1996c), for grain yield and other traits (distinct genetic controls), a breeder should realize that the performance of untested single crosses can be effectively predicted using relationship information from molecular or pedigree data, unbalanced and large data set, and diverse heterotic patterns. The significance of genomic prediction has been confirmed with maize (Zhao et al. 2015) and other important crops, as rice (Xu et al. 2014), wheat (Zhao et al. 2013b) and barley (Philipp et al. 2016), along the last 10 years. Why, then, is there no published evidence that prediction of untested single crosses is of general use by breeders of worldwide seed companies? What should be additionally proved to make prediction of untested single crosses as successful as the Jenkins’ (1934) method for predicting double crosses performance was? We believe that this paper offers a significant contribution.

Our assessment on efficiency of prediction of untested single cross performance keeps some similarities with few earlier studies but sharp differences for most previous investigations. This study is based on simulated data set, as the study of Technow et al. (2012), assuming 400 QTLs distributed along ten chromosomes. Thus, the prediction accuracies and coincidence indexes (a measure of untested single crosses selection efficiency) are available for non-assessed single crosses since the values were computed based on the true genotypic values of the non-assessed single crosses and not on a cross-validation procedure involving assessed single crosses. This does not mean that we consider simulated data better than field data or have any criticism on the cross-validation procedure. We know that simulated data, because the assumptions, cannot integrally describe the complexity of populations and genetic determination of traits (Daetwyler et al. 2013). To highlight the relevance of (overall) LD, our study is based on scenarios not favorable to prediction of untested single cross performance: very low level of relationship between the DH lines, low and intermediate heritabilities for the assessed single crosses, and not higher heterotic pattern. In the studies of Massman et al. (2013) and Bernardo (1994, 1995, 1996a) the relationship among inbreds from the same heterotic group ranged from 0.11 to 0.58. Riedelsheimer et al. (2012) observed high relationship only between the non-Stiff Stalk inbreds. Technow et al. (2012) assumed non-related inbreds. For most of the investigations on prediction of untested single crosses and testcrosses, the grain yield heritability ranged from 0.72 to 0.88. The common heterotic patterns in these previous studies are Stiff Stalk and non-Stiff Stalk, and Dent and Flint. The MAF in the groups of Dent and Flint inbreds were approximately 0.10 and 0.20, respectively, and approximately 20% of the SNPs showed a difference of allelic frequency of at least 0.6.

Concerning the prediction accuracy and the efficiency of identification of the superior 300 non-assessed single crosses, our results prove that prediction of untested single crosses is a very efficient procedure (note that we are not saying genomic prediction), especially for low and intermediate heritabilities of the assessed single crosses. The prediction accuracy of the non-assessed single crosses under low (0.55 to 0.71) and intermediate (0.74 to 0.87) accuracies of assessed single crosses achieved 0.85 and 0.89, respectively. It is important to highlight that these are not relative accuracies. Most important, the coincidence of the non-assessed single crosses under low (0.26 to 0.39) and intermediate (0.44 to 0.66) parametric coincidences of assessed single crosses achieved 0.59 and 0.64, respectively. For high heritability (80 to 95%; accuracies from 0.89 to 0.97), as observed in most of the studies on prediction of untested single cross performance, we can state (based on values predicted by fitting a quadratic regression model) that the prediction accuracy of non-assessed single crosses is up to only 10% lower (0.87 to 0.92) and, most impressive, the coincidence index can range from 0.61 to 0.71 (parametric coincidences between 0.72 to 0.93). Under maximum accuracy of assessed single crosses (1.0), the prediction accuracy and coincidence of non-assessed single crosses achieved 0.93 and 0.76. Thus, assuming high heritability, high density, and training set size of 30%, the accuracy can achieve 0.92 and the efficiency of identification of the best 9% of the non-assessed single crosses can achieve 0.71. It is important to highlight that this efficacy can be higher by using more related DH or inbred lines, under high LD. Thus, we strong recommend that maize breeders, as well as rice, wheat, and barley breeders, make widespread use of prediction of non-assessed single crosses, at least for preliminary screening or prior to field testing.

To take advantage of genomic prediction, Kadam et al. (2016) recommend redesigning hybrid breeding programs. However, because breeders are unlikely to rely solely on genomic predictions when selecting superior untested hybrids, Technow et al. (2014) believe that genomic prediction will be combined with field testing of the most promising experimental hybrids. For grain yield, the prediction accuracies observed by Bernardo (1994, 1995, 1996a) ranged from 0.14 to 0.80, proportional to the heritability (in the range 35–74%) and training set size. The non-relative accuracies (relative accuracy x root square of heritability) observed in the studies of Kadam et al. (2016), Technow et al. (2014), Massman et al. (2013), Technow et al. (2012), and Riedelsheimer et al. (2012) ranged between 0.20 and 0.86, also proportional to the heritability (in the range 53–98%) and training set size.

We hope that readers of this paper have realized the importance of (overall) LD for effective prediction of non-assessed single crosses, as well as genetic variability (see the parametric accuracy of genomic prediction). Breeders have no control over LD and relatedness between the DH or inbred lines. However, selection should always provide high level of overall LD in the groups of selected DH or inbred lines. Comparison of our LD assessment with the LD analyses from other studies is inadequate because we have distances in cM and not in base-pairs. But in general the level of LD was high (r^2^ of approximately 0.3) only for SNPs separated by up to 0.5 Mb (Technow et al. 2014; Massman et al. 2013; Technow et al. 2012; Riedelsheimer et al. 2012). To maximize the prediction accuracy and the efficiency of identification of the best non-assessed single crosses it is necessary to adopt the random sampling of single crosses for testing instead of the random sampling of DH or inbred lines for a diallel. This is because sampling 30 or even 10% of the single crosses leads to single crosses for testing derived from all DH or inbred lines from each group. In our case, in every resampling assuming training set size of 30 and 10% we always get groups of assessed single crosses (1,470 and 490 single crosses, respectively) derived from the 70 DH lines of each group. However, sampling DH lines for a diallel provided 1,440 and 484 single crosses for testing derived from 38 and 22 DH lines, respectively. Thus, the sampling of single crosses provides best prediction of the SNP average effects of substitution. Riedelsheimer et al. (2012) emphasized the need for large genetic variability to obtain high prediction accuracies. Further, their results indicated that pairs of closely related lines and population structuring only weakly contributed to the high prediction accuracies. Regarding dominance, because it can be a relevant genetic effect, breeders should always fit the additive-dominance model to maximize the prediction accuracy and the efficiency of identification of the best non-assessed single crosses. Interestingly, in most of the studies on prediction of non-assessed single crosses the prediction accuracy did not significantly increase when modeling SCA in addition to GCA effects (Zhao et al. 2015).

Concerning SNP density and training set size, factors related with the costs of genotyping and phenotyping, breeders should find a balance between efficiency and expenses, since maximizing SNP density and training set size maximizes the efficiency of untested single cross prediction. Based on our results, because the decreases in the prediction accuracy (approximately 4%) and coincidence index (5 to 10%) by decreasing the average SNP density from 0.1 to 1 cM are of reduced magnitude, we consider sufficient to employ custom genotyping to provide an average SNP density of 1 cM. Decreasing the training set size from 30 to 10% of the single crosses does not significantly affect the prediction accuracy under intermediate to high heritability (decrease of up to 9%), but the coincidence index can be reduced in up to 21%. However, considering that the coincidence index will be kept in the range 0.48 to 0.61, proportional to the heritability, and that the maximum values are in the range 0.48 to 0.61, we also consider sufficient to assess at least 10% of the possible single crosses. As highlighted by Zhao et al. (2015), marker density only marginally affects the prediction accuracy of untested single crosses and, for biparental populations, a plateau for the accuracy is reached with a few hundred markers. Technow et al. (2014) did not find an improvement of prediction accuracies by using higher SNP density. Additionally, the increase in the training set size led to a relative small increase in the prediction accuracy. However, the prediction accuracies obtained by Riedelsheimer et al. (2012) under high density (38,019 SNPs) were substantially greater than those reached with a low-density marker panel (1,152 SNPs). In the study of Technow et al. (2012), the prediction accuracies increased with SNP density and number of parents tested in hybrid combination.

The DH lines sampling process, the heterotic pattern, and the statistical approach should not be worries for breeders. However, under high heritability notice that sampling more than one DH line per S_0_ or S_3_ plant provided the higher coincidence values and high prediction accuracy in our study. For rice, wheat, and barley breeders our message is: high prediction accuracy and high efficiency of identification of superior non-assessed single crosses does not depend on heterotic groups but on the (overall) LD in the group or in each group of DH or inbred lines. In other words, the efficiency of prediction of non-assessed single crosses derived from DH or inbred lines from the same population can be as high as the efficiency of prediction of untested single crosses derived from DH or inbred lines from distinct heterotic groups. This is not confirmed comparing the relative prediction accuracies for grain yield of maize untested single crosses (from approximately 0.50 to 0.95, for most studies) with those obtained with rice, wheat, and barley untested hybrids (0.50 to 0.60, approximately) (Philipp et al. 2016; Xu et al. 2014; Zhao et al. 2013b). However, the lower relative prediction accuracies for untested rice, wheat, and barley hybrids should be due to prediction of two- and three-way crosses. Regarding the statistical approach, our model did not provide an increase in the efficiency of non-assessed single cross prediction, compared to the models proposed by Massman et al. (2013) and Technow et al. (2012). It is important to highlight that our results showed that these two models are really identical (data no shown). Thus, because the simplified definition of the incidence matrices for these two previous models, it is quite safe to use any of them. Finally, the choice between the statistical approaches RR-BLUP (prediction of genotypic values of non-assessed single crosses based on prediction of SNP average effects of substitution), GBLUP (prediction of genotypic values of non-assessed single crosses based on additive and dominance genomic matrices), and BLUP (prediction of genotypic values of non-assessed single crosses based on additive and dominance matrices from pedigree records) is not a serious worry for breeders too. Our evidence is that there is no significant difference between RR-BLUP and GBLUP regarding prediction accuracy and efficiency of identification of the best untested single crosses. Further, even when the level of relatedness between the DH or inbred lines in each group is low, in general pedigree-based BLUP is as efficient as genomic prediction, excepting when the DH lines are derived from inbred population. Thus, DNA polymorphism is not essential for an efficient prediction of non-assessed single cross performance. In a review on genomic selection in hybrid breeding, Zhao et al. (2015) state that the choice of the biometrical model has no substantial impact on the prediction accuracy of untested single crosses. Technow et al. (2014) observed that prediction methods GBLUP and BayesB resulted in very similar prediction accuracies. According to Massman et al. (2013), pedigree-based BLUP and RR-BLUP models did not lead to prediction accuracies that differed significantly. Comparing GBLUP and BayesB, Technow et al. (2012) concluded that the latter method produced significantly higher accuracies for the additive-dominance model.

Our main contributions on the assessment of prediction efficiency of untested single cross performance are: 1) the prediction accuracy of untested single crosses ranged from approximately 0.80 to 0.90 as the heritability of tested single crosses ranged from low (30%) to high (100%); however, the efficacy of identification of the best 9% of the untested single crosses ranged from approximately 0.50 to 0.70, depending on the DH lines sampling process; 2) the prediction accuracy for crops showing no defined heterotic pattern can be as efficient as with maize, for which there are well defined heterotic groups; this is because the most important factor affecting the prediction efficiency is the overall LD; 3) to maximize prediction accuracy and coincidence the choice of single crosses for testing should be based on a random process; this procedure maximizes the number of DH lines in hybrid combinations and provides better predictions of the SNP average effects of substitution and dominance deviations, compared to sampling DH lines for a diallel; 4) because non significant decreases in the prediction accuracy and coincidence, the prediction of untested single crosses can be efficient assuming reduced training set size (10%) and SNP density of 1 cM; 5) RR-BLUP and GBLUP provide equivalent prediction efficiencies of untested single crosses; 6) excepting for DH lines derived from inbred populations, pedigree-based BLUP is as efficient as genomic prediction of untested single crosses; and 7) the theoretical accuracy shows that the prediction accuracy is not affected by the linkage phase.

## ACKNOWLEDGMENTS

We thank the National Council for Scientific and Technological Development (CNPq), the Brazilian Federal Agency for Support and Evaluation of Graduate Education (Capes) and the Foundation for Research Support of Minas Gerais State (Fapemig) for financial support.

